# Plasminogen repairs abnormal pain perception through improving sensory function recovery and regeneration of peripheral small nerve fiber in db/db mice

**DOI:** 10.1101/792325

**Authors:** Weiquan Li, Ting Wang, Fen Chen, Chunying Guo, Yanghui Liao, Congcong Quan, Fei Zheng, Jinan Li

**Author notes:** Corresponding author, Room C602G, 289 Digital Peninsula, Shunfeng Industrial Park, No.2 Red Willow Road, Futian District, Shenzhen, China *E-mail address* (J.N. Li).

## Abstract

Painful diabetic peripheral neuropathy (PDPN) is a devastating complication of diabetes and severely threatens the health of humankind. The plasminogen activator system and plasminogen (Plg) have multiple functional roles in tissue regeneration and extracellular matrix remodeling, which suggests that Plg may have a potentially pivotal role in anti-PDPN. In the present study, we explore whether an increased level of circulating Plg has positive effect on repairing abnormal pain perception in diabetic mice model. Our data demonstrated that additional Plg not only helps healing pain allodynia or hyperalgesia on the mice at the age of 8 weeks old in early PDPN, but more important, also has positive effects of regaining normal pain perception from hypoalgesia on the mice at ages of 14-15 or 24-25 weeks in advanced PDPN. Furthermore, our data also reveal a possible mechanism for Plg’s contribution to rebuilding normal pain perception among db/db mice by promoting axonal myelination and regeneration of small nerve fiber in peripheral nervous system. Therefore, our data suggest that Plg show promise to become a drug candidate for treating diabetic peripheral neuropathic pain.

## Introduction

Painful diabetic peripheral neuropathy (PDPN) is a common complication accompanying long term Diabetes Mellitus (DM), affecting approximately 50% diabetic patients [1]. PDPN has been recently defined as a symmetric, length-dependent sensorimotor polyneuropathy attributable to metabolic and microvascular alterations as a result of chronic hyperglycemia exposure [2]. Though the specific pathogenesis of PDPN in different stages has not been fully clarified, but it is known that there are two stages according to the manifestations, the early PDPN and the advanced PDPN. Such manifestations are 1) thermal hyperalgesia, an equivalent of a clinical phenomenon described in early PDPN; 2) thermal hypoalgesia, typically present in advanced PDPN; 3) mechanical hyperalgesia, an equivalent of pain on pressure in early PDPN; 4) mechanical hypoalgesia, an equivalent to the loss of sensitivity to mechanical noxious stimuli in advanced PDPN 5) tactile allodynia, a painful perception of a light touch [3]. 20-30% of patients with PDPN suffer from severe neuropathic pain [4–5], as a leading cause for foot ulceration and amputation and fall related injury. This may result in withdrawal from social events there by affecting the quality of life and considerably increasing the financial burden of treatment [6–8].

Unfortunately, so far, there are no effective FDA approved drugs available for treating PDPN [9–10]. At present, traditional drugs including (1) antiepileptic drugs, such as gabapentin and pregabalin; (2) analgesics and anesthetics; (3) antidepressants and non-steroidal anti-inflammatory drugs have been used to treat diabetic neuropathic pain [9–10]. However, only a small portion of PDPN patients (about less than 20%) shows response to these treatments. Furthermore, the high side effects, high cost and insufficient effects of these drugs have limited their use in treating PDPN. Therefore, there is an urgent need for the research and development of effective medications to relieve PDPN.

Plasminogen (Plg) is a zymogen mainly produced by the liver and activated to become the broad-spectrum protease plasmin by either of two physiological plasminogen activators (PAs): tissue-type PA (tPA) or urokinase-type PA (uPA) [11]. It has been shown that the extracellular proteolytic activity of plasmin plays a pivotal role in fibrinolysis and extracellular matrix degradation, which is essential during many damaged tissue remodeling processes, including diabetic wound healing [12–14]. Furthermore, our previous study showed that administering Plg improved diabetic wound healing by tissue remodelling [14]. Interestingly the research on mice from Seeds’ group has demonstrated that deficiency of any components in the PA system, such as Plg or tPA, will delay sensational response to external stimuli [15]. However, the roles of Plg on PDPN is an emerging area.

In order to make sure the role of Plg in PDPN, a diabetic (db/db) mouse model was applied to dissect the effects and function of Plg in PDPN and to explore related underlying mechanism. Our findings support that Plg may play a critical role in alleviating pain allodynia through rebuilding normal pain perception by significantly promoting repair and regeneration of damaged small somatosensory nerve fiber. Therefore, Plg may be a promising therapeutic candidate in treating PDPN and preventing its related diseases.

## Materials and methods

### Animals

Leptin receptor-deficient db/db mice and their littermates were obtained from the Animal Research Center of Nanjing University. The animals were kept under standard laboratory conditions. The Ethics Committee of Talengen Institute of Life Sciences approved all of the experimental protocols. For studying diabetic peripheral neuropathic pain, the male animals with age at 8, 14-15 and 25-26 weeks were grouped and treated with or without Plg protein for indicated time depending on the experimental design. In the experiments of a burn-wound model, the 16-26 weeks old db/db male mice were grouped and treated with Plg for the experiments. If not mentioned, at least five mice were included in each experimental group.

### h-Plg protein administration in diabetic burn-wound healing and diabetic peripheral neuropathy study

For the study of diabetic burn-wound healing, the male mice at age 16-26 weeks old were anesthetized by an intraperitoneal injection of 50 mg/Kg sodium pentobarbital. A copper rod was heated to 95-100°C by submersion in boiling water. The copper rod was immediately applied vertically for 6 seconds without additional pressure on the back skin of mice that had also been depilated before wounding. After wounding, all mice were individually caged, and wounds were neither sutured nor dressed. The mice received standardized wounds, and then 2 mg of human Plg was administered daily by 0.2ml IV injection. In the control group, 0.2 ml of PBS was administered daily by IV injection as a placebo. The daily treatments were continued for indicated days depending on the experimental design.

### h-Plg protein administration in diabetic peripheral neuropathy study

As for the study of diabetic peripheral neuropathy, each group of male mice at the age of 8 weeks, 14-15 and 24-25 weeks were divided into two subgroups, and 2 mg of human Plg daily was administered directly on Plg-treated group mice by 0.2 ml IV injection, and the control group mice were administered 0.2 ml of PBS daily by IV injection as a placebo. The daily treatments were continued for indicated days depending on the experimental design. Then the animals were used for designed behavior testing.

### Behavior quantitative sensory testing (QST)

Behavioral signs representing three different components of neuropathic pain were examined: allodynia, hyperalgesia, and hypoalgesia in response to cold and mechanical stimuli. To quantify mechanical sensitivity of the foot, the standard quantitative sensory testing (QST) was used to record numbers of brisk foot withdrawal in response to normally noxious and innocuous mechanical stimuli as described previously [16–17]. Von Frey fibers are a neurophysiological examination tool used for determining the mechanical pain threshold in humans and nonhumans [17]. The force applied to the testing animal is based on the size of monofilament in von Frey used. First, an intermediate size of monofilament (number 4.31, exerting 2.0 g of force) was gently applied with enough force to bend it. This was repeated up to three times in distinct areas along the lateral paw. In the case of a positive response (rapid withdrawal of the paw within 2 seconds), a smaller filament was tested. If no response was recorded in any of the three different areas, a larger filament was tested. The frequency of foot withdrawal was expressed as a percent: (# of trials accompanied by brisk foot withdrawal) × 100 / (# of total trials). On a given test day and for each hind paw, the same procedure was repeated by using two different sizes of von Frey filaments. Mechanical sensitivity was tested on each mouse on day 0 (1 day before administration of Plg) and 3, 4, 6, 7, 9, 11, 12 and 16 days after treating with Plg.

To quantify cold sensitivity of the foot, brisk foot withdrawal in response to acetone application was measured. The mice were placed under a transparent plastic dome on a metal mesh floor, and acetone was applied to the plantar surface of the foot. To do this, an acetone bubble was formed at the end of a small polyethylene tube that was connected to a syringe. The bubble was then gently touched to the heel. The acetone quickly spread over the proximal half of the plantar surface of the foot. The acetone was applied five times (once every 5 min) to each paw. The frequency of foot withdrawal was again expressed as a percent: (# of trials accompanied by brisk foot withdrawal) × 100 / (# of total trials). Cold sensitivity was tested on each mouse on day 0 (1 day before administration of Plg) and 3, 4, 6, 7, 11, 12, and 16 days after treating with Plg.

To quantify the pain sensation evoked by pin-prick, the 27-gauge needle was used to stimulate the foot of the mouse but not to penetrate the dermis. The db/db mice were stimulated on the soles of their left and right feet every 3 minutes for a total of 10 times, and the number of paw withdrawal reactions was counted. The frequency of foot withdrawal was again expressed as a percent: (# of trials accompanied by brisk foot withdrawal) × 100 / (# of total trials). Pin-prick test was performed on each mouse on day 0 (1 day before administration of Plg) and 3, 4, 6, 7, 11, 12, and 16 days after treating with Plg.

### H&E staining

The sciatic nerve tissues were fixed in 4% paraformaldehyde, embedded in paraffin and sectioned 3 μm thick. The sections were stained for morphological analysis using an H&E staining kit. The slides were examined by light microscopy under a Nikon microscope, and images were recorded digitally using a camera connected to a computer.

### Immunohistochemical analyses

The paraffin-embedded sections were rehydrated and then treated with antibodies against Fibrin (cat# ab27913), or PGP9.5 (cat# ab10404) purchased from Abcam (Cambridge, UK). The signal intensity was detected immunohistochemically by the peroxidase anti-peroxidase method. In brief, the antigens were first retrieved by treatment with Citrate buffer at high temperature for 20 min, and then the tissue sections were blocked with 5% non-immunized goat serum (Vector laboratories, USA, cat# SK-1012-50) and incubated with the antibodies against Fibrin or PGP9.5 diluted in PBS. After this procedure, an anti-rabbit link antibody was applied, followed by a rabbit PAP complex. The staining was visualized through a diaminobenzidine (DAB) reaction, and the sections were counterstained with hematoxylin.

### Light microscopic examination

The slides were examined by light microscopy under a Nikon microscope (C-SHG1), and images were recorded digitally using Nikon DS-Fi3 connected to a computer. When needed, IPP6.0 pathological image analysis software was used to determine the integral optical density (IOD) or average optical density (AOD) of immuno-positive products for each group and area of interest. The integrated optical density (IOD) is the sum of the individual pixels within the field of view of the specimen. Average optical density (AOD) is the total value of pixels in the area of interest (IOD SUM) divided by the area to be tested. Both IOD and AOD represent the intensity of protein expression.

## Results

### Effect of Plg on small sensory nerve fiber in wounds sites in diabetic mice

To explore the effect of Plg on peripheral nerve fiber in diabetic wound sites, the diabetic db/db mice were treated daily for 3, 7 or 14 days by intravenous (IV) injections of 2 mg/0.2 ml human Plg or 0.2 ml PBS after burning injury. PGP9.5 immunohistochemical staining of a tissue section from wounded sites was performed. The results (Fig.1A-B) showed that the expression of PGP9.5 in the wound sites of the Plg-treated group is higher than the control group without Plg administration on day 3 and 7, and with a significantly difference on day 14 (p<0.05). The results indicated that Plg treatment enhances the regeneration of injured small nerve fiber while the healing of diabetic wounds was improved.

### Effect of Plg on sciatic nerve fiber in the diabetic mice

To further investigate the effect of Plg on sciatic nerve fiber in the diabetic mice, the db/db mice at the age of 24-25 weeks old were injected intravenously with 2 mg Plg protein daily for 15 days. The H&E staining of sciatic nerve tissue shows that a large number of the myelin sheath of axons are swollen, demyelinated and disintegrated in the control group without administration of Plg (as shown in figure 2A(a)). In contrast, sciatic nerve fibers of Plg-treated mice were morphologically closely arranged, and the myelin sheath of the axonal structure was well maintained, and few of them are disintegrated as shown in figure 2A(b). Moreover, increased expression of PGP9.5 of sciatic nerve tissue in the Plg-treated group was observed by PGP9.5 immunohistochemical staining test (Fig. 2B). These data indicate that the number and density of sciatic nerve fibers with intact axonal structures are increased and better maintained in the Plg-treated group.

Increased deposition of fibrin at the injured site is a common feature of diabetic injury. In turn, the reduction of fibrin deposition may contribute to the healing of injured tissue. To investigate the effect of Plg on fibrin in wound sciatic nerve tissue, the 24-25 weeks old db/db mice were injected intravenously with 2 mg Plg daily for 15 days. Then the samples of injured sciatic nerve tissues from these mice were subjected to detect the degree of fibrin deposition by fibrin immunohistochemical staining test. As shown in figure 2C, the deposition of fibrin in sciatic nerve tissue in the Plg-treated group was significantly lower compared to the control group. This result suggests that increased Plg in circulation facilitates fibrin degradation to clear its deposition, which may be beneficial for axonal regeneration in damaged sciatic nerve tissue of db/db mouse model.

### Effect of Plg on both hyperalgesia and hypoalgesia to pain sensation in diabetes mice

The db/db mice develop diabetes at 4 weeks of age, exhibit allodynia and hyperalgesia between 8 and 12 weeks of age, and exhibit hypoalgesia after 12 weeks of age [18]. In the present study, we followed a protocol to quantify the degree of mechanical allodynia in response to pressure from a von-Frey filament. From Fig.3A, the results showed that the pain threshold of the group from 8 week old mice administrated with Plg after 9 days is significantly increased (P< 0.05), indicating the symptom of diabetic hyperalgesia is improved.

In contrast, the results from 14-15 week old mice with advanced PDPN representing hypoalgesia showed that the pain threshold of the group administrated with Plg gradually reduced and exhibited significant difference on days 3 and 12 compared with the control group without Plg treatment (Fig. 3B, P<0.05). Similarly, the results from 24-25 week old mice with advanced PDPN showed that the pain threshold of the group administrated with Plg gradually reduced and exhibited significant difference on day 16 (Fig. 3C, P<0.05). Together, Plg can increase the pain threshold of diabetes mice at early PDPN and decrease the pain threshold of diabetes mice at advanced PDPN.

### Effect of Plg on abnormal thermal perception to cold stimulation in diabetic mice

Damaged peripheral small sensory nerve systems due to diabetic hyperglycemia can also change the thermal perception resulting in cold allodynia in response to cold stimuli. To explore if Plg treatment has effects on thermal perception to cold stimuli, two groups of diabetic mice treated with Plg at different ages were subjected to cold stimulation behavior test with acetone. The results of the QST from the Plg-treated 14-15 week old mice showed that a significant increase of sensational response to cold stimulation on days 6 (P<0.05)and 12 (P<0.01) compared with the control group without Plg treatment (Fig. 4A). Similarly, the results from the Plg-treated 24-25 week old mice show a significantly increased percentage of sensational response on days 4 and 16 of treatment (Fig. 4B). It should also be noted that in all days with non-significant results, the mean values of the treated group still trend in the same direction as the significant ones. Our results indicate that Plg treatment may repair diabetic-induced somatosensory dysfunction to relieve pain and cold allodynia in db/db mouse model with advanced PDPN.

### Effect of Plg on abnormal pain sensation evoked by pinprick test in diabetic mice

To further investigate the effect of Plg on somatosensory nervous system in the diabetic mice model, two groups of diabetic mice treated with Plg protein at different ages were subjected to pin-pricking behavior tests. The results from the Plg-treated 14-15 week old mice showed that the percentage of sensational response to pin-prick test was significantly increased on days 6 (P<0.05)and 12 (P<0.001)compared with the control group without Plg treatment (Fig. 5A). Similarly, the results from the Plg-treated 24-25 week old mice with advanced PDPN showed a significant increase of sensational response on days 7 (P<0.01) and 16 (P<0.05) (Fig. 5B). Just as mentioned above, the mean values of the treated group on all non-significant days trend to the same direction as the significant ones. The results indicate that Plg treatment may alleviate pain hypoalgesia in the diabetic mouse model with advanced PDPN.

## Discussion

Painful diabetes peripheral neuropathy (PDPN) is a subtype of sensorimotor neuropathy and is the most commonly acquired neuropathy in diabetes mellitus (DM) [19–20]. Its pathomechanism has not yet been completely clarified, but the neuronal hyperexcitability and sensitization in early stage of DPN and hypalgesia in late stage of DPN are widely accepted as playing an important role.

Allodynia and hyperalgesia at early stage of DPN characterized by spontaneous or evoked pain is described as electric shock-like or burning after nerve dysfunction and damage in the peripheral sensory nerve system induced by diabetic hyperglycemia. In the peripheral nervous system (PNS), there are more thin (<1 μm) unmyelinated axons, known as C-fiber axons or small fibers, than myelinated axons [21]. Therefore, the earliest changes of DPN occur at the level of unmyelinated C fibers, with initial imbalance between degeneration and regeneration of C fiber resulting in pain, allodynia, and hyperalgesias [22]. In the present study, we observed that Plg treatment is able to correct abnormal pain sensitization by increasing the pain threshold on the db/db mice at early stage of PDPN after 9 days of Plg supplementation when symptom of hyperalgesia, exhibiting hypersensitive to mechanic pressure, is developed (Fig. 3A). It is known that the impairment of small nerve fibers results in the loss of normal thermal and pain perception, whereas large nerve fiber impairment results in loss of normal touch and vibration perception [23–25]. Small nerve fibers constitute 70 to 90% of peripheral nerve fibers and are functionally classified into somatic sensory, somatic motor, and autonomic fibers, which regulate several key functions such as tissue blood flow, temperature, and pain perception as well as sweating [26–27]. Expression of PGP9.5 in the diabetic wound skin is increased after administration with Plg for 14 days (Fig.1). This results indicated that Plg promoted small nerve fibers regeneration at the wounded site in the diabetic mice. Blasi and Mignatti have disclosed that the PA system was implicated in various tissue remodelling processes as early as 1990s’ [28–29]. That is to say Plg can improve allodynia and hyperalgesia of early stage of PDN through promoting the regeneration of small nerve fibers to elevate the pain perception threshold.

As the disease course progresses, mild segmental axonal demyelination then occurs, followed by frank axonal degeneration of myelinated fibers as demyelination surpasses remyelination[30]. These changes lead to a progressive loss of distal sensation in a distal-to proximal course along the nerve that defines diabetic peripheral neuropathy [1]. This phase is defined as the advanced stage of DPN. At this stage, hypoalgesia is developed and exhibiting hyposensitive to external mechanic, cold stimuli and mechanical noxious stimuli. Surprisingly, we observed that Plg treatment could reverse pain insensate state to normal pain response by lower the pain perception threshold of diabetic mice at the age of 14-15 weeks and 24-25 weeks that has lost sensation to the pain and developed advanced DPN (Fig.3B-C, Fig.4, and Fig.5). Advanced DPN is associated with elevated vibration and thermal perception thresholds that progress to sensory nerve loss, occurring in conjunction with degeneration of all fiber types [3]. In the current study, H&E staining result shown that the nerve fibers in sciatic nerve of diabetic mice administrated with Plg were wrapped by epineurium and closely arranged, while a large number of myelin axons in sciatic nerve tissue of diabetic mice without Plg treatment are swelling, demyelinated and collapsed (Fig. 2A). Meanwhile, increased expression of PGP9.5 in sciatic nerve tissue of Plg-treated group indicated that the number and density of small nerve fibers with intact axonal structures are increased and better maintained in the Plg-treated group (Fig. 2B). In addition, the deposition of fibrin in sciatic nerve tissue of the Plg-treated group was significantly decrease (Fig.2C). These results revealed that Plg can improve hypoalgesia by promoting sciatic nerve remyelination of diabetic mice to lower sensory nerve damage and loss,

The pathogenesis of DPN is considered to be multifactorial. A combination of chronic hyperglycemia, oxidative stress, inflammation, microangiopathy of nerve blood vessels and a reduction in nerve fiber repair result in damage to the vasa nervorum, the microvessels that supply blood to neural tissue[31–32]. Chronic hyperglycaemia and hyperlipidemia exposure in diabetes mellitus cause mitochondrial dysfunction and inflammatory signals activation, further leading to metabolic injury and cell apoptosis, ultimately resulting in microvascular and neuronal damage. The unmyelinated C fibers are more susceptible to metabolic injury in large part because they lack the degree of protection and nutrient supplementation afforded to myelinated axons by Schwann cells (SCs) [33]. Furthermore, high glucose increases ROS and inflammatory factor level through the TLR4/NFκB pathway in Schwann cells (SCs) [34]. All of these will cause the peripheral nerve cell death and demyelination as lacking nutrient supplementation of SCs and microvascular. Recently, Juliana’ group revealed that Plg/Pla and the receptor Plg-R_KT_ could mediate macrophage polarization to the M2 phenotype via STAT3 signaling, playing a key role in anti-inflammatory [35]. Meanwhile, PA system/plasminogen is reported to regulate expression and activation of some growth factors including transforming growth factor (TGF-β), nerve growth factor (NGF), vascular endothelial cell growth factor (VEGF) and fibroblast growth factor (FGF), and other growth factors for tissue regeneration and extracellular matrix remodeling [36]. Further studies revealed that activated TGF-β stimulates Schwann cell proliferation and differentiation which is essential for the formation of axonal myelin sheaths and axonal support for nerve regeneration and functional recovery in the peripheral nervous system after injury [37–38]. Zou’s study have showed that tPA or tPA/Plg promotes axonal remyelination and regeneration of injured sciatic nerve tissue[39]. Therefore, Plg can promote peripheral nervous axons remyelination and regeneration by multiple pathway to repair allodynia and hyperalgesia and reverse hypoalgesia of PDPN in diabetes mellitus.

## Acknowledgments

This work was supported by grants from the National Natural Science Foundation of China (Grant No.81570741) and Guangdong Provincial Department of Science and Technology, China (Grant No. 2014A020210001).

## Figure legends

**Fig. 1. Plg treatment promotes the regeneration of small nerve fiber in diabetic wounds**

A: Representative image of PGP9.5 immunohistochemical staining of wounded skin of db/db mice on days 3, 7, 14 after injury respectively. (a-c): Control group, (d-f): Plg-treated group, Magnification is 400x.

B: Quantitative analysis of PGP9.5 expression in figure 1A. n = 5 mice used in each group. * P < 0.05 vs. Control group.

**Fig. 2. Plg treatment promotes regeneration of injured sciatic nerve in the diabetic mice**

**A:** Representative image of H&E staining of sciatic nerve tissue from db/db mice at the age of 24-25 weeks treated with Plg for 15 days, (a): Control group, (b): Plg-treated group, Magnification is 400x.

B: Representative image of PGP.95 immunohistochemical staining of injured sciatic nerve of db/db mice at age of 24-25 weeks treated with Plg for 15 days, (a): Control group, (b): Plg-treated group, Magnification is 400x.

C: Representative image of fibrin immunohistochemical staining of injured sciatic nerve tissue of db/db mice at the age of 24-25 weeks treated with Plg for 15 days, (a): Control group, (b): Plg-treated group, Magnification is 400x.

**Fig. 3. Plg treatment alleviates mechanical allodynia by behavioral von-Frey filament test of diabetic mice**

A: Quantitative analysis of mechanical allodynia and hyperalgesia on eight-week-old db/db mice with early-stage DPN treated by Plg supplementation for the indicated number of days.

B: Quantitative analysis of mechanical allodynia and hypoalgesia on 14-15 week old db/db mice with middle stage of DPN treated by Plg supplementation for the indicated number of days.

C: Quantitative analysis of mechanical allodynia and hypoalgesia on 24-25 week old db/db mice with advanced DPN treated by Plg supplementation for the indicated number of days.

**Fig. 4. Plg treatment enhances sensational response to cold stimulation by acetone in diabetic mice**

A: Quantitative analysis of cold allodynia and hypoalgesia on 14-15 week old diabetic mice treated by Plg supplementation for the indicated number of days.

B: Quantitative analysis of cold allodynia and hypoalgesia on 24-25 week old diabetic mice treated by Plg supplementation for the indicated number of days.

**Fig. 5. Plg treatment enhances sensational response to pain stimulated by pin-prick test in diabetic mice**

A: Quantitative analysis of mechanical allodynia and hypoalgesia on 14-15 week old diabetic mice treated by Plg supplementation for the indicated number of days.

B: Quantitative analysis of mechanical allodynia and hypoalgesia on 24-25 week old diabetic mice treated by Plg supplementation for the indicated number of days.

